## Supplementary Information for "Simultaneous polyclonal antibody sequencing and epitope mapping by cryo electron microscopy and mass spectrometry – a perspective"


**Contents:**


- Legend to Supplementary Tables S1-2
- Supplementary Figures S1-2

**Supplementary Table S1.** Benchmark of experimental cryoEM maps of monoclonal antibody-antigen complexes from EMDB.

*Name*: PDB-ID of deposited model for corresponding map

*Actual*: corresponding to true VH/VL sequence as deposited in PDB model

*Built:* corresponding to *de novo* sequences determined by ModelAngelo

*Segment*: inferred V-gene from top-scoring hit in Stitch alignment

*Score:* alignment score in Stitch

*Identity*: sequence identity between *Actual-Built* based on *de novo* seq. or inferred *Segments*

*Resolution:* Global FSC resolution of corresponding map

*HC*: heavy chain

*LC:* light chain

**Supplementary Table S2.** Benchmark of EMPEM maps downloaded from EMDB for *de novo* modelling in ModelAngelo, with alignment scores in Stitch.

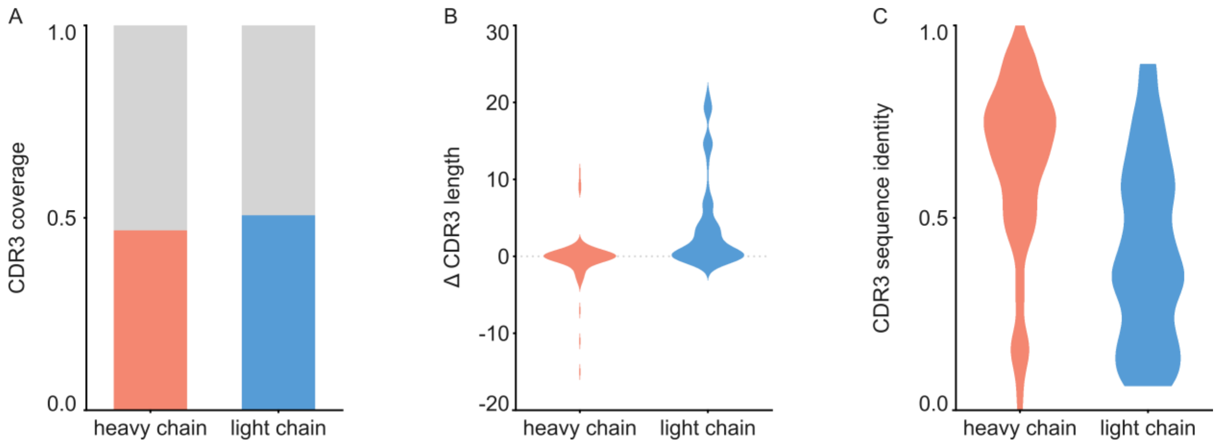


**Supplementary Figure S1.** Analysis of *de novo* CDR3 modelling in ModelAngelo-Stitch. A) Coverage of CDR3 for the heavy and light chain. CDR3 was counted for coverage if the *de novo* sequence spanned the flanking cysteine on the V gene and the tryptophan or phenylalanine on the J gene. Proportion of maps with CDR3 coverage in red/blue, maps with missing CDR3 in grey. B) Difference in length between *de novo* modelled CDR3 vs. true sequence. C) Sequence identity of *de novo* modelled CDR3 vs. true sequence

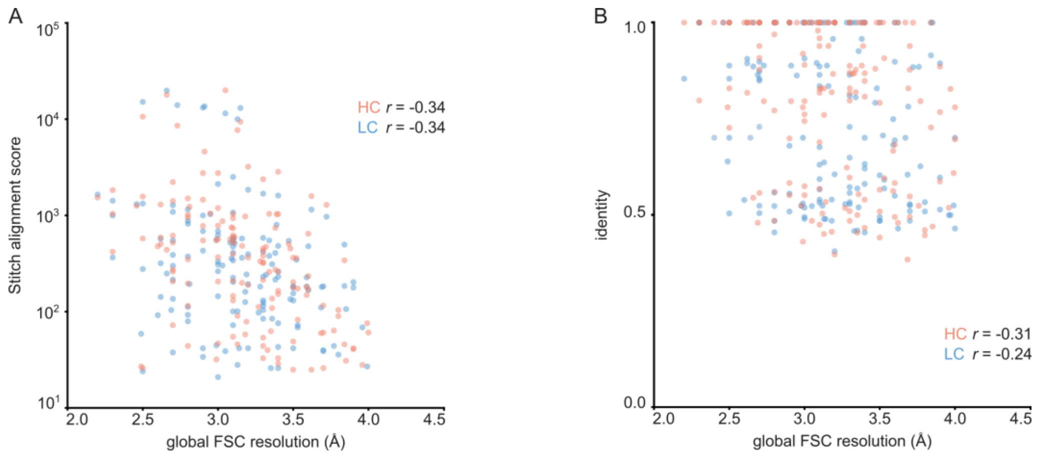


**Supplementary Figure S2.** Correlation between global FSC resolution and Stitch alignment score (A) or inferred V-gene identity. The non-parametric Spearman correlation coefficient is indicated for heavy and light chain.
